# An open source platform for presenting dynamic visual stimuli

**DOI:** 10.1101/2020.12.24.424344

**Authors:** Kyra Swanson, Samantha R. White, Michael W. Preston, Joshua Wilson, Meagan Mitchell, Mark Laubach

## Abstract

Operant behavior procedures often rely on visual stimuli to cue the initiation or secession of a response, and to provide a means for discriminating between two or more simultaneously available responses. While primate and human studies typically use LCD or OLED monitors and touch screens, rodent studies use a variety of methods to present visual cues ranging from traditional incandescent light bulbs, single LEDs, and, more recently, touch screen monitors. Commercially available systems for visual stimulus presentation are costly, challenging to customize, and are typically closed source. We developed an open-source, highly-modifiable visual stimulus presentation platform that can be combined with a 3D-printed operant response device. The device uses an eight by eight matrix of LEDs, and can be expanded to control much larger LED matrices. Implementing the platform is low-cost (<$70 USD per device in the year 2020). Using the platform, we trained rats to make nosepoke responses and discriminate between two distinct visual cues in a location-independent manner. This visual stimulus presentation platform is a cost-effective way to implement complex visually-guided operant behavior, including the use of moving or dynamically changing visual stimuli.

**Significance Statement:** The design of an open source and low cost device for presenting visual stimuli is described. It is capable of presenting complex visual patterns and dynamically changing stimuli. A practical demonstration of the device is also reported, from an experiment in which rats performed a luminance based visual discrimination. The device has utility for studying visual processing, psychophysics, and decision making in a variety of species.

## Introduction

Visual stimuli are a crucial component for studying behavioral neuroscience. Studies of perception, cognition, learning, memory, decision making, among many others, often rely on the presentation of visual cues to probe response behaviors from multiple species. The varieties of visual stimuli that have been implemented in research studies over the last few decades range from simple to complex. In studies of rodents utilizing visual cues, subjects are often trained to respond to the presence or location of illumination as in Pavlovian conditioning procedures (e.g. Saunders et al., 2018), the five-choice serial reaction-time task (Bari et al., 2008), and decision making tasks (e.g. Raposo et al., 2012). While these studies are complex, the hardware necessary is as simple as a single lightbulb or LED. However, the visual cues therefore lack the flexibility to manipulate multiple parameters of visual stimuli, especially in real-time. To address this issue, several labs have adopted the use of commercial LCD screens or touch screen systems to present more complex visual stimuli such as contrast gratings, textured shapes, naturalistic cues, or 3D rendered objects (Busse et al., 2011; Vinken et al., 2014; De Keyser et al., 2015; Djurdjevic et al., 2018). These screens contain more pixels per square inch and are capable of displaying more detailed images, but require a means to store the images in memory (or require a computer to run software for displaying images) and are rather expensive. These displays may also require significant reconfiguration of a behavioral chamber for optimal placement or the purchase of dedicated chambers for testing with video screens and the use of the screens can limit the spatial locations where cues are presented on the experimental participants.

To expand the complexity of visual cues while preserving flexibility in the behavioral set up, we developed a simple but highly customizable platform for visual stimulus presentation utilizing affordable and readily available parts. The device reported here is distinct from several previous papers that have reported using Arduino and single LEDs or LED matrices for visual stimulus presentation (Pinto et al., 2011; Teikari et al., 2012; Jones et al., 2016). Our open-source platform consists of three LED matrices that can be independently controlled via an open-source microcontroller (Arduino Uno) to display a variety of complex visual stimuli. The matrices are held within a 3D printed frame which can be easily mounted at various locations in a behavioral chamber, including above nose poke ports, lever presses, or reward spouts. This system can be flexibly integrated with established behavior set-ups as it only requires TTL serial inputs from any existing set-up to the microcontroller. Here, we provide instructions and share the necessary resources for building and customizing the visual display system which can be made for under $70 USD (Fall 2020). We demonstrate validity for use of this platform by implementing it in a behavioral task to examine perceptual decision making in rats.

## METHODS

### LED Matrices

Three Pure Green 1.2” 8×8 LED matrices (Adafruit) were used for visual stimulus presentation. We note that other LED matrices and graphics control libraries are available, such as the NeoPixel system (Adafruit), and the software can be modified to support these devices. The matrices use an inter-integrated communication (I^2^C or I2C) protocol and require a microcontroller ‘primary’ device that is I2C-capable, such as an Arduino, to control the ‘secondary’ matrix devices (https://learn.sparkfun.com/tutorials/i2c/all). The I2C protocol is vulnerable to electrical noise in long bus cables, so we advise that the microcontroller be kept in close proximity to the matrices with short bus cables. The LED matrices are screwed into 3D-printed trays that vertically orient the matrices (Extended Data; GitHub). The trays can then be glued to custom-made 3D-printed nosepoke ports (optional - Extended Data; GitHub), or for a semi-permanent option, can be mounted with velcro.

These devices are controlled using the open source Adafruit_GFX and Adafruit_LEDBackpack libraries. We provide microcontroller software (Extended Data; GitHub) that is preconfigured to switch between 16 programmed visual stimulus sets (see Extended Data or Github for details). Each of the 16 possible stimuli is mapped to a unique combination of the binary voltage status of the four digital input pins on the Arduino. (An alternative implementation using the teensy microcontroller platform was also developed. Files and instructions for using this version of the device are available upon request from the authors.) The digital pins on the Arudino need to be grounded via one of the GND pins off the Arduino and a common ground from the source of the device that sets the voltage on the pins to high or low. This is achieved by using a breadboard with 10k pull down resistors for each digital pin (Figure 1). The source of V+ inputs can vary by experimental set up; in our experimental set up, MedPC SmartControl sends a 28V output to toggle a relay, sending a 5V+ signal to the digital pins on the Arduino (see Extended Data or GitHub for specifics about integrating MedPC Smart Control outputs with an Arduino via a relay).

**Figure 1:**
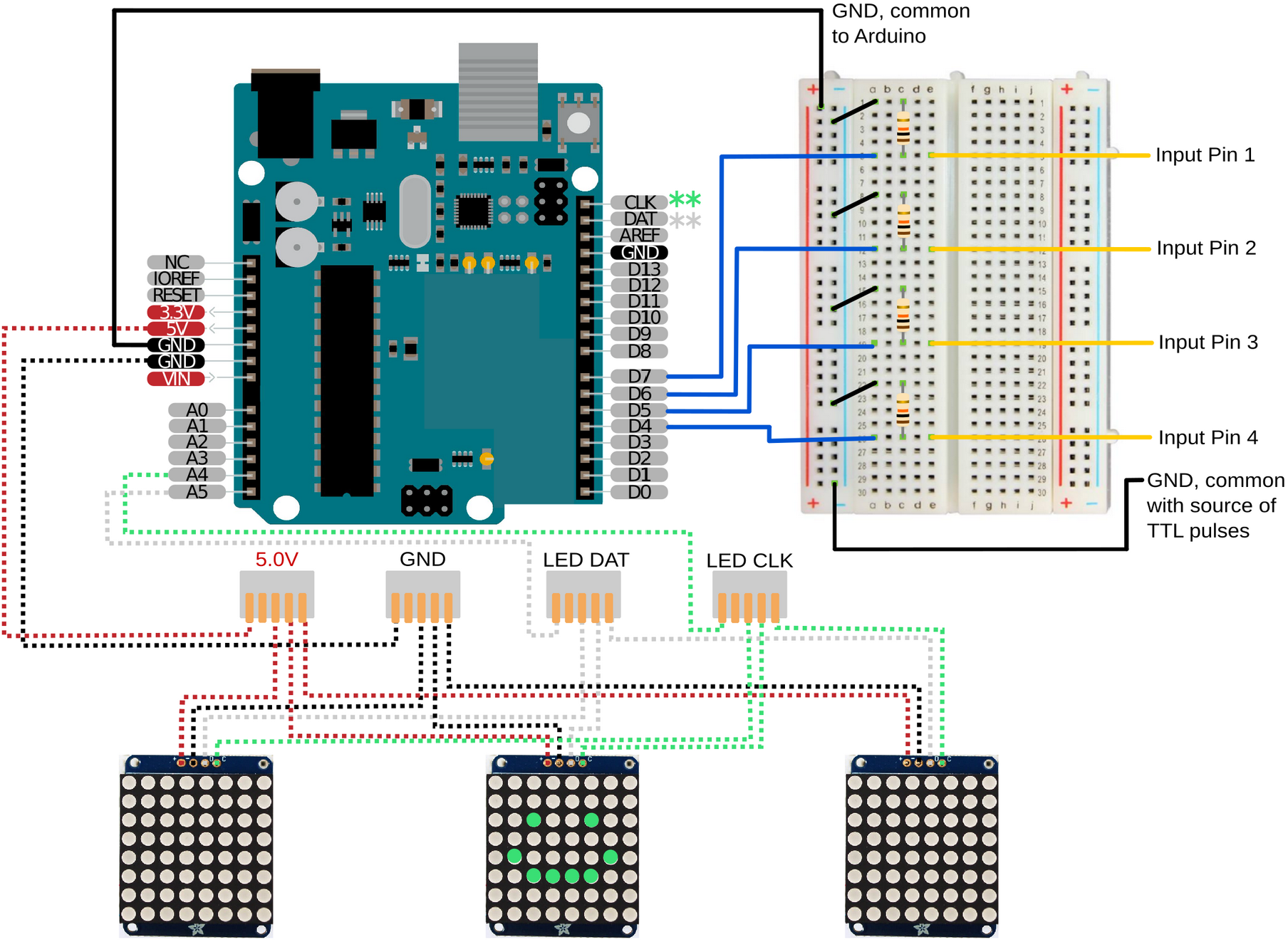
Hardware and wiring for the LED matrices. The required wiring is shown for connecting the Arduino microcontroller (upper left) to the LED matrices (lower row) and the extenal control system that selects stimulus patterns based on TTL serial inputs. The version reported here allows for stimulus control by four distinct TTL inputs. Connectivity with the external control system can easily be arranged using a standard breakout board (upper right). Pins on the backpack controller for LED matrices are shown above the LED matrices.

We programmed the software for use with three individual LED matrices. This way, the matrices can be placed next to three separate operant responses. Each input combination turns on a distinct stimulus on one or multiple LED matrices, which we call a stimulus set. The stimulus set can indicate which of three operant responses, in this case nosepokes at specific positions, will correspond to which outcomes on each trial. The stimulus set can indicate, for example, that only one operant response is active and available, or that the animal must choose between multiple responses with the same or different outcomes (Figure 2A). The pin combination-stimulus mapping is modifiable, as is the number of input pins and corresponding number of stimuli. The software can support up to 2^n^ distinct stimuli, where *n* is the number of available input pins on the microcontroller. While the software currently supports 3 matrices, this can also be changed. I2C can support as few as one operative device, up to the available number of unique device addresses. The 1.2” 8×8 matrix used here supports up to 8 addresses.

**Figure 2:**
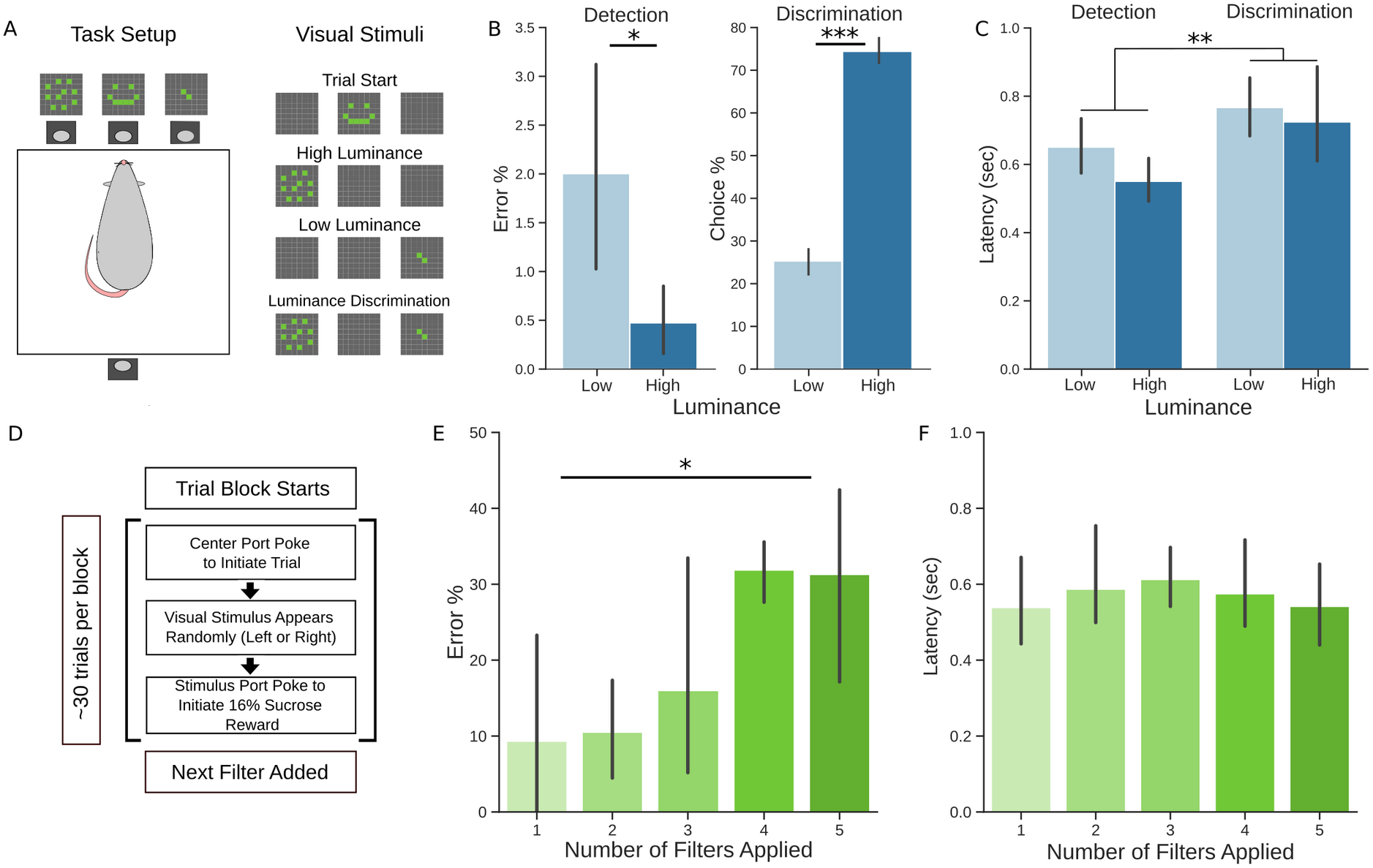
Visual discrimination behavior using the LED matrices. A) Rats were trained and tested in an operant chamber with the LED matrices mounted above response ports on one side of the chamber and with a reward port containing a spout mounted on the opposite side of the chamber. Each trial was initiated by a nosepoke response in the center port to the “Trial Start” cue, followed by the presentation of either the high or low luminance cue. Both cues were presented for discrimination trials. B) In detection trials, rats expressed a greater number of errors when detecting the low luminance cue compared to the high luminance cue. In discrimination trials, rats chose the high luminance cue more often than the low luminance cue. C) Rats responded more slowly on discrimination trials compared to detection trials. D) Reduced luminance testing: The test session started with one filter layer over the LED matrices and rats had to report the low luminance cue on the left or right nosepoke port. After approximately every 30 trials, additional filter layers were added for a total of five layers. E) As each layer was added, the overall error percentage increased. F) Response latencies were stable over the level of luminance. Asterisks: * = p<0.05; ** = p<0.01; *** = p<0.001. Error bars represent 95% confidence intervals.

The luminance of the LED matrices over a range of illuminated pixles was measured using a Thorlabs PM100D, set to the frequency range of the green LEDs. The measured values are reported in Table 1.

**Table 1:** Luminance levels for the LED matrices with 1 to 16 pixels illuminated.

| Number of LED Pixels Illuminated | PeakLuminance (in $\mu\text{W}$) |
| --- | --- |
| 1 | 6.2 |
| 2 | 9.3 |
| 4 | 18.5 |
| 8 | 30.2 |
| 16 | 58.0 |

### Behavioral Apparatus

All animals were trained in sound-attenuating behavioral boxes (ENV-018MD-EMS: Med Associates). A single horizontally placed spout (5/16” sipper tube: Ancare) was mounted to a lickometer (MedAssociates) on one wall, 6.5 cm from the floor and a single green LED light was placed 4 cm above the spout (henceforth referred to as the spout light). The opposite wall had three of the 3D-printed nosepoke ports aligned horizontally 5 cm from the floor and 4 cm apart, with the IR beam break sensors on the external side of the wall. The nosepoke ports contain slots designed for Adafruit 3mm IR Break Beam sensors, which function at 3.3 or 5V. The slots make it possible to easily insert and remove the sensors as needed. Additionally, the nosepoke ports feature an 8.4mm diameter centered hole within the recessed portion of the port. This can be used to deliver fluid or odors to the nosepoke port. Finally, the front of the nosepoke port contains 4 holes for mounting screws and a cutout for a cap which can be used to temporarily block entry to the nosepoke port for training purposes. Three Pure Green 1.2” 8×8 LED matrices were used for visual stimulus presentation and were placed 2.5 cm above the center of each nosepoke port, outside the box. Data collection and behavioral devices, including the Arduino that drove the LED matrices, were controlled using custom-written code for the MedPC system, version IV (Med Associates).

### Subjects

Twenty-four male Long–Evans rats (300–450 g, Charles-River) were individually housed and kept on a 12 h light dark cycle with lights on at 7:00 AM. Facilities were temperature and humidity controlled. Rats were given several days to acclimate to the facilities, during which they were handled and had free access to food and water. During training and testing, animals were on regulated access to food to maintain their body weights at approximately 90% of their free-access weights. All animal procedures were approved by the American University Institutional Animal Care and Use Committee (Washington, DC).

### Training Procedure

Animals were first trained to lick at a reward spout to receive ~50 μL of a 16% wt/vol liquid sucrose in the presence of a visual stimulus (5 mm green LED), located above the reward port. Over the next several sessions, animals were hand-shaped to respond in nosepoke ports in response to distinct visual stimuli to initiate trials, report the location of visual stimuli, and gain access to liquid sucrose rewards. A cue (4×4 square of illuminated pixels) was displayed over the center port for trial initiation, which was then followed by a high luminance (8 illuminated pixels) or low luminance (2 illuminated pixels) cue, randomized by side and luminance intensity. Responses at the port below the high luminance cue response yielded access to a 16% wt/vol sucrose reward at the reward spout, while responses at the port below the low luminance cue response yielded access to a 4% wt/vol sucrose reward. When animals performed at least ~120 trials in a 60 minute session during the training phase, they were moved to testing detection/discrimination testing.

### Detection/Discrimination Testing

Twenty of the animals were tested in this procedure. As in training, they initiated trials by nosepoking at the center port. For detection trials (~67% of total trials), animals were presented with a single stimulus randomized by side and luminance intensity. For discrimination trials (the remaining ~33% of trials), animals were presented with both the high and low luminance cue randomized by side. The rats always received 16% sucrose at the reward port after they responded in the port below the high luminance stimulus and always received 4% sucrose after they responded in the port below the low luminance stimulus. They received no fluid if they responded below a non-illuminated port on nonchoice trials. We measured the ‘response latency’ as the time taken for rats to go from trial initiation port to the illuminated choice port after the onset of the visual stimuli. We also measured the number of detection errors and the response rate for each cue when both stimuli were presented. Rats did not show improved discrimination/choice behavior over the period of discrimination testing, as the percentage of choices for the higher and lower value stimuli was relatively stable over the period of testing. They did, however, show reduced differences in choice latencies for the higher and lower value stimuli.

### Luminance Testing

Four additional rats were tested to determine their ability to detect 2 illuminated pixels under decreasing luminance. These animals completed initial training, were not tested using the detection and discrimination procedures described above, and were tested as described here.

Luminance was controlled by placing transparent layers of film (Rosco E-Colour #210.6 Neutral Density) over the LED matrices. In single test sessions, filters were added, one layer at a time every 28±6 trials up to five layers in total. The filters reduced luminance from 9.0μW to <0.1 μW with five layers (Table 2). We measured response latencies as previously described as well as the number of detection errors for each level of luminance.

**Table 2:** Luminance levels for 2 illuminated pixels with 0 to 5 filters placed over the LED matrices.

| Number of Filters over 2 LED pixels | Peak Luminance (in $\mu\text{W}$) |
| --- | --- |
| 0 | 9.3 |
| 1 | 2.5 |
| 2 | 0.7 |
| 3 | 0.2 |
| 4 | 0.1 |
| 5 | $< 0.1$ |

### Data Analysis

Behavioral data were saved in standard MedPC data files and were analyzed using custom-written code in the Python and R languages. *Response latency* was defined as the time elapsed from the initiation of a trial (prompted by a center port response) to the nosepoke response in the left or right port. Response latencies greater than 2 seconds were filtered out of the data to exclude trials where rats disengaged from the task. For the remaining response latencies, the median response latency was calculated per rat. Oneway ANOVAs were performed in R to examine effects of luminance and trial type (discrimination vs detection trials) on response latency for the Detection/Discrimination experiment or examine the effect of number of filters on response latency in the Luminance experiment.

*Errors* in detection trials occurred where rats produced a nosepoke response in a non-illuminated port. Error percentages were calculated by dividing the number of error trials by the total number of detection trials per rat. *Choice percentage* in discrimination trials reflects the rate at which rats responded to either the high luminance or low luminance target. These were calculated by dividing the number of high or low luminance trials by the total number of discrimination trials per rat. Paired t tests and one-way ANOVAs were performed in R (*aov* function) to examine effects of luminance and trial type (discrimination vs detection trials) on error percentage or choice percentage for the Detection/Discrimination experiment or examine the effect of number of filters on error/choice percentage in the Luminance experiment.

### Build Instructions

All the software code, 3D print files can be found in Extended Data and on GitHub. Briefly, start by uploading the software onto the microcontroller; ensure that the identity of the digital input pins assigned in the code match your desired input pins on the Arduino (in our example we used digital pins 4-7). Next, the LED matrices must be soldered to the included backpack and uniquely addressed. Instructions for this process are on <u>Adafruit’s website</u>. Our software code identifies three unique addresses (x70, x71, and x74); the code can easily be modified to include different addresses. Next, cables to connect the 5V, GND, SCL and SDA pins on the microcontroller must be soldered to each of the respective pins on the matrices (Figure 1); alternatively, you can create a solderless connector by creating a jump wire to attach to the pins on the backpacks. We recommend the use of a 5-way wire connector (Figure 1) when connecting the respective wires between the backpacks and the microcontroller to increase flexibility in the number of backpacks that can be implemented in future iterations. Mount the matrices onto the 3D-printed trays and secure to the nosepoke ports with either glue or velcro. Place the microcontroller keeping the bus cable as short as possible. Then connect the 5V+ TTL output pins to the designated row of the breadboard for the desired input pins (D4-D7 in our set up). We recommend using a prototyping shield and screw-down terminals to secure the wires for maximum stability. Supply power to the device and test matrix function. Inconsistent flickering or incorrect display will likely be due to loose connections on the breadboard, misaddressed backpacks, misaddressed input pins on the Arduino, or bus cables that are too long.

### Code Accessibility

The software and hardware design files described in the paper are freely available online at [URL redacted for double-blind review]. The files are available as Extended Data. The software was tested on Intel and AMD based personal computers running the Microsoft Windows 10 operating system and the Linux Mint 20 Cinnamon operating system.

## RESULTS

### Detection/Discrimination Testing

To assess whether rats could reliably detect and discriminate cues, we presented cues of high or low luminance one at a time (detection) or both at the same time (discrimination trials) (Figure 2A). We calculated their error and choice percentages for high and low luminance cues. In detection trials, rats expressed a greater number of errors when detecting the low luminance cue compared to the high luminance cue (Figure 2B, left; F(1,37)=7.192, p<0.05). n discrimination trials, rats chose the high luminance cue more often than the low luminance cue (Figure 2B, right; t(19)=-16.305, p<0.001). We also measured their response latencies across trial types and cue luminance to assess if there was an effect of luminance on latency. Overall, rats expressed a greater latency in discrimination trials compared to detection trials (Figure 2C; F(1,76)=9.674, p<0.01) with no detectable difference due to luminance in this population of rats (F(1,76)=2.342, p=0.13011). Together these data suggest that the cues are both detectable and discriminable to rats in a basic decision making task.

It may be surprising to some readers that rats choose the low luminance target ~25% of the time, Reinforcement with 4% liquid sucrose was provided at the reward port when animals responded at the choice port below the low luminance stimulus. This concentration of liquid sucrose has been shown to maintain responding in a number of previous studies (e.g. Flaherty et al. 1983; Barr and Phillips, 1999; Taha and Fields, 2005; Parent et al., 2015). A detailed analysis of the effects of varying sucrose concentration on multiple measures of consummatory behavior found that hungry rats licked more and showed higher instrumental breakpoints when working for 4% sucrose compared to plain water in a progressive ratio licking task (Swanson et al., 2019). Beyond having value as a nutrient, our animals were forced to respond for this amount of sucrose when presented with the lower luminance stimulus over many hundreds of trials during training. There were no negative consequences of choosing the lower luminance stimulus other than receiving a lower concentration of sucrose at the reward port. For these reasons, the fraction of choice trials with choices of the lower value target might be higher than in other studies, e.g. in classic studies of visual discrimination learning and two alternative forced choice designs that typically use reward vs no reward.

### Luminance Testing

To assess the limits of detectability, a subset of four rats completed a series of low luminance detection trials. The test session started with one filter layer over the LED matrices and rats were expected to detect the cue over the left or right choice port. After approximately thirty trials (28±6), an additional filter layer was added (Figure 2E). The session completed after four additional filter layers were added, for a total of five layers. We calculated the error percentages for each block of trials where each additional filter layer was added. We found that the overall error percentage increased as the number of filters increased (Figure 2E; F(4,13)=4.119, p<0.05). We also measured response latencies and found no significant increase or decrease in median latency as a function of filter layers (Figure 2F; F(4,13)=0.194, p=0.937). Together these data suggest that the cues are suitable for psychophysics testing in rodents.

## DISCUSSION

We present a versatile, highly-modifiable visual stimulus presentation platform for behavioral research. This platform uses a standard Arduino Uno microcontroller board and LED matrices from Adafruit. It is open-source and low-cost. It provides up to 16 unique stimului across three separate LED matrices. We provide instructions for making the device, a bill of materials for all required parts, and extended data including example programs for using the device. We also provide feasibility data based on psychophysical testing of Long-Evans rats performing a visual discrimination task using the device.

Our device differs in several important ways from existing devices that are commonly used in visual testing, especially for rodents. First, the device is based on an 8 by 8 matrix of LEDs and not an LCD monitor. LCD monitors have been used in a number of classic (Bussey et al., 1997) and more recent studies (Zoccolan et al., 2009; Zoccolan, 2015; Broschard et al., 2019; Nikbakht and Diamond, 2021), and low cost Arduino compatible LCD screen are available (e.g. http://www.dsdtech-global.com/) that could be used as an alternative to the LED matrices from Adafruit used in the present device. LCD monitors have an advantage for presenting highly detailed visual stimuli, beyond the 8×8 “pixels” available in the Adafruit LED matrices. LCD screens are capable of producing more than one color and can be rapidly changed for studies such as gradient discrimination (e.g. Zoccolan et al., 2009). We note that the ability to vary color may not matter much for studies of rodent vision, as reported here, given that rats and mice only perceive colors in the green and ultraviolet ranges (e.g. Jacobs et al., 2001).

Second, LCD screens have limited viewing angles and generally slower temporal responses compared to LEDs due to the liquid crystal reactions that underlie the functioning of an LCD screen. These issues are not practical problems for the kind of visual discrimination study reported here, but could matter if the effects of gaze position is relevant to the brain area of interest (Meister, 2018) or if one is interested in the precise timing of a neuronal response in the visual pathway of a behaving animal (e.g. Durand et al., 2016).

Third, our device does not include a touch sensitive screen, which has become common in behavioral testing in rodents in recent years. Such a screen could easily be added to the device, if one desires, perhaps using components from an open source touch screen system (MouseBytes: https://www.mousebytes.ca/).

Our visual stimulus presentation platform is highly flexible and customizable. The 64-pixel LED matrices are capable of delivering a variety of visual cues, from single pixels to complex shapes, including moving patterns of stimuli. They are small, relatively cheap, and come in several colors. The microcontroller software can switch between 16 preset visual cues across three individual LED matrices. An example of an interesting use of the device, not reported here, would be to change the presented stimuli during the time required for animals to perceive differences in the stimuli or make a decision and examine processes such as “conflict” about the stimuli. By modifying the example code provided in this report, it is possible to create a number of preset cues for many different kinds of behavioral experiments. An additional, perhaps overlooked benefit of using a microcontroller rather than a hardwired commercial LED device is that updated software can be loaded quickly and often, even between sessions or cohorts on the same day.

Furthermore, the LED panels have three dimensions that make them versatile as visual stimuli. The location of the LED matrices within the box, the brightness of the LEDs when on, and the number and positions of illuminated LEDs within the matrix all can be manipulated to create distinct visual cues. RGB matrices can add color as a fourth dimension, assuming the colors chosen are detectable by the animal model. For example, contrasting Green and White stimuli in rodent behavioral testing can add another level of information. Or one could make custom matrices using other ranges of color, such as ultraviolet, which would be especially interesting for the study of visual discrimination in rodents.

The mechanism for attaching the LED mount to the nosepoke port ensures quick removal and replacement when necessary, while guaranteeing stability of location of the LED matrix day to day. Importantly, the entire device is low-cost, modifiable, and easily cleaned and replaced. The 3D-printed nosepoke ports have many useful features, especially for early-stage behavioral training. The cap temporarily but completely blocks access to a nosepoke port, without the need to detach the port from the behavioral chamber. The additional hole within the recession can be used to deliver fluid or odor cues. The mount orients the matrix to be perpendicular to the ground and places it directly above the nosepoke port. We have found this design to be especially useful for training approach behavior in rats during the initial period of training.

Using this platform, we trained rats to accurately discriminate between two visual cues. The rats showed increased errors in detecting the lower luminance stimuli compared to higher luminance stimuli and higher probabilities of choosing the brighter stimulus (Figure 2B). These effects were also apparent in their choice latencies (Figure 2C). Using sets of filters that reduced the intensity of the stimuli (Table 2), we carried out detection testing in a subset of rats and found that errors in detection varied with luminance (Figure 2E), without any overall effect on detection latency (Figure 2F).

Classic studies of rodent vision reported higher accuracies than reported here (Lashley, 1930). In those classic studies, the rats experienced strongly negative outcomes for making an incorrect choice (falling from a platform into a net). By contrast, our rats received positive outcomes for responses to both the higher and lower value stimuli. These differences in the behavioral design might account for the differences in choices of the correct stimuli in Lashley’s studies and of the higher value stimulus in ours.

The device reported here is a low cost and flexible option for conducting visual experiments in rodents and other species. It readily can be integrated into existing behavioral control systems that use TTL serial communication. The provided programs can be modified to allow for more complex patterns of stimuli and for dynamic changes in stimuli during the experiment. The opensource nature of the design can be modified by any user based on their own research questions and does not depend on approval or input from any commercial developer or vendor (White et al., 2019). All component parts are provided in the Parts List. We hope that our design is useful to the community and will allow for new approaches to evaluating visual processing in a variety of species.

### Extended Data

The following files are included in the Extended Data:

- VisualStimuli-Arduino-eNeuro.ino – Arduino code for the device.
- LED-Matrix-Holder-Part1.stl and LED-Matrix-Holder-Part2.stl – Design files for the 3D printed holders for the LED matrices.
- nosepoke.stl and nosepoke-block.stl – Design files for the optional nosepoke ports.
- FullSetUp.png – Image showing wiring diagram for LED Backpacks, Arduino, breadboard, and relays to integrate with MedPC control system.
- WiringDiagram.png – Image showing specific wiring for the LED Backpacks, Arduno, and breadboard.
- LEDMatrix-Addressing.png – Documentation on addressing for the LED Backpacks.
- StimCodes.png – Documentation on producing the visual patterns over three LED matrices as described in this manuscript.
- README.txt – Overview on files in the Extended Data, and available on project’s GitHub page.

## Extended Data / GitHub Repository

https://github.com/LaubachLab/LED-matrices

## Author contributions

KS, MWP, and ML developed the initial hardware design for the Arduino platform; KS extended the design to the teensy platform; SW revised the design for Arduino, as reported here; KS and MWP wrote the software; KS and JW designed 3D printed parts; SW, MM, and ML carried out the behavioral experiments; SW analyzed the behavioral data; KS, SW and ML wrote the paper.

## Acknowledgements

We thank Dr Wade Kothman for helpful discussion on the filters used for luminance testing.

## Parts List

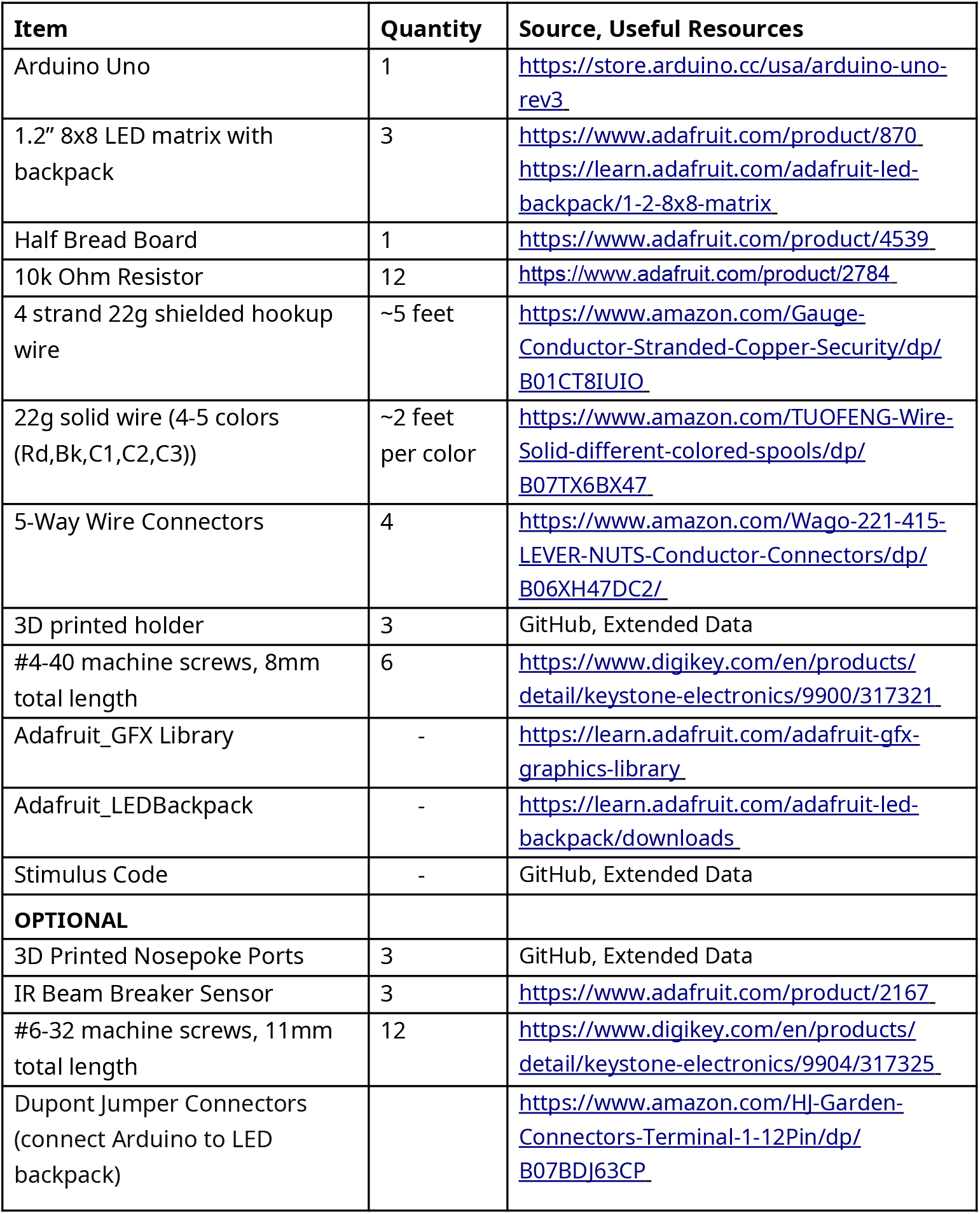

| Item | Quantity | Source, Useful Resources |
| --- | --- | --- |
| Arduino Uno | 1 | https://store.arduino.cc/usa/arduino-uno-rev3 |
| 1.2" 8x8 LED matrix with backpack | 3 | https://www.adafruit.com/product/870<br>https://learn.adafruit.com/adafruit-led-backpack/1-2-8x8-matrix |
| Half Bread Board | 1 | https://www.adafruit.com/product/4539 |
| 10k Ohm Resistor | 12 | https://www.adafruit.com/product/2784 |
| 4 strand 22g shielded hookup wire | ~5 feet | https://www.amazon.com/Gauge-Conductor-Stranded-Copper-Security/dp/B01CT8IUIO |
| 22g solid wire (4-5 colors (Rd,Bk,C1,C2,C3)) | ~2 feet per color | https://www.amazon.com/TUOFENG-Wire-Solid-different-colored-spools/dp/B07TX6BX47 |
| 5-Way Wire Connectors | 4 | https://www.amazon.com/Wago-221-415-LEVER-NUTS-Conductor-Connectors/dp/B06XH47DC2/ |
| 3D printed holder | 3 | GitHub, Extended Data |
| #4-40 machine screws, 8mm total length | 6 | https://www.digikey.com/en/products/detail/keystone-electronics/9900/317321 |
| Adafruit_GFX Library | - | https://learn.adafruit.com/adafruit-gfx-graphics-library |
| Adafruit_LEDBackpack | - | https://learn.adafruit.com/adafruit-led-backpack/downloads |
| Stimulus Code | - | GitHub, Extended Data |
| **OPTIONAL** |  |  |
| 3D Printed Nosepoke Ports | 3 | GitHub, Extended Data |
| IR Beam Breaker Sensor | 3 | https://www.adafruit.com/product/2167 |
| #6-32 machine screws, 11mm total length | 12 | https://www.digikey.com/en/products/detail/keystone-electronics/9904/317325 |
| Dupont Jumper Connectors (connect Arduino to LED backpack) |  | https://www.amazon.com/HJ-Garden-Connectors-Terminal-1-12Pin/dp/B07BDJ63CP |

